# Multi-omics analyses of single cell-derived colorectal cancer organoids reveal intratumor heterogeneity and immune response diversity

**DOI:** 10.1101/2022.11.25.517889

**Authors:** Jian-Hui Yue, Jie Li, Qu-Miao Xu, Qi-Wang Ma, Chao Chen, Song-Ming Liu, Hai-Xi Sun, Qiao-Qi Sui, Feng Mu, Pei-Rong Ding, Long-Qi Liu, Mirna Perez-Moreno, Xi Zhang

## Abstract

Colorectal cancer (CRC) organoids have similar genomic and functional characteristics to the original specimens. They have become a novel and promising tumor model. However, systematic research of heterogeneity and evolution based on tumor organoids and unbiased evaluation of CD4^+^ T cell-mediated tumor recognition is presently lacking. Here, we study the variability in genomic characterization and single-cell transcriptome between different tumor organoid clones derived from the same CRC patient. While clone-specific differences in driver gene mutations and cancer stem cell (CSC) subpopulations were observed, inter-clone heterogeneity was even more prevalent. Using single-cell RNA sequencing, we found that compared with bulk organoids, CRC organoid clones are mainly composed of cancer stem cells and transiently-amplifying cells (TACs), but no epithelial-mesenchymal transition (EMT)-like (S100A4^+^CDH1^-^) cells. In addition, we study the phenotypic characteristics and TCR properties of tumor-reactive CD4^+^ T cells by coupling the organoid-TILs co-culture model, single-cell profiling, and recognition testing of candidate TCRs. We found that the proliferating CD4^+^ T cells increased significantly from 7.1% to 35.4%, and the TCR clonotypes increased from 110 to 1043 after co-culture with organoids. Moreover, 4 of 10 (40%) candidate TCRs with a significant increase in clone abundance exhibited tumor recognition. These results provide evidence that the proliferating CD4^+^ T cells with clonal expansion after co-culture with tumor organoids are the potential tumor-reactive T cells. Collectively, we reconstructed intra-patient heterogeneity in single cell-derived organoid models and characterized the phenotypic characteristics and functional properties of tumor-reactive CD4^+^ T cells.

## Introduction

Based on the statistics from GLOBOCAN 2020, colorectal cancer has become the second most deadly cancer worldwide in 2020(*1*). The overall survival of most CRC remains poor due to late diagnosis, resistance to chemotherapy or immunotherapy, and tumor metastasis(*1, 2*). Cancer evolution and heterogeneity are the main intrinsic factors leading to cancer progression and resistance to treatments. Even individual cancer cells derived from a single cancer biopsy can independently acquire specific genetic and epigenetic characteristics, and with increased heterogeneity, they displayed significantly divergent responses to chemotherapy(*3-5*). Reconstructing intra-patient heterogeneity in novel and promising tumor models is essential for understanding tumorigenesis and tumor progression and provides valuable resources to develop personalized immunotherapies.

Patient-derived tumor organoids (PDTOs) are three-dimensional cellular structures derived from tumor-specific stem cells which recapitulate the genomic and functional resemblances features of the original specimens(*5-7*). Currently, a few living organoid biobanks of CRC patients have been established and widely used to predict the response to chemotherapy immunotherapy, and to study the tumor microenvironment(*8-10*). Some recent studies showed that single cell-derived tumor organoids from CRC patients displayed diversity in HLA class I peptide presentation and responses to small-molecule chemo-drugs, making the organoid clones an ideal study system intra-tumor heterogeneity(*5, 11*). Since single cell-derived organoids might originate from different cancer stem cells, they could facilitate the study of tumor clonal heterogeneity, such as clonal evolution, functional diversity, or identification of low-frequency tumor mutations. So far, systematic studies of CRC evolution and heterogeneity in genetic, epigenetic, and microenvironment based on single cell-derived organoids are still lacking.

Besides intrinsic cancer factors, the tumor microenvironment provides the “soil” for tumor cells to grow, where cancer evolution is largely shaped by immune editing. Tumor-infiltrating lymphocytes (TILs) are vital in tumor control; however, a large number of studies have shown that TILs acquire exhaustive phenotypes in the tumor microenvironment during chronic exposure to tumor antigens. Expansion of TILs from cancer patients *in vitro* restores effector functions, and clinical studies demonstrated that TIL therapies effectively treat patients with metastatic melanoma, colorectal cancer, and lung cancer. Notably, as cytotoxic CD8^+^ T cells are well known to eradicate tumors, recent studies highlighted that cytotoxic CD4^+^ T cells induced strong antitumor immune responses in mouse models and human bladder cancer patients(*12-14*). Adoptive transfer of autologous tumor-mutation-specific CD4^+^ TILs mediated effective tumor regression in a patient with metastatic cholangiocarcinoma(*15*). Some results suggested that CD39, CD137, PD-1, CD103 are potential markers to identify tumor-reactive CD8+ T cells(*16-18*). However, little is known about the phenotypic characteristics and functional properties of tumor-reactive CD4^+^ T cells in the malignant tumor, hindering the development of immunotherapies based on cytotoxic CD4^+^ T cells.

Here, we generated single cell-derived organoids from CRC patients and demonstrated that they display CRC intra-tumor heterogeneity at the genetic and transcriptomic levels. Furthermore, by co-culturing CRC bulk organoids with TILs, we studied the phenotypic characteristics and functional properties of tumor-reactive CD4^+^ T cells through single-cell profiling and tumor recognition testing of candidate T cells receptors (TCRs). We demonstrated that the clonal expansion of proliferating CD4^+^ T cells leads to tumor-reactive T cell clonotypes, which might provide useful TCR sequences for developing personalized TCR-engineered cell therapies against tumors. These findings will ultimately help improve personalized immunotherapies targeting tumor stem cells and further understand the CD4^+^ T cells mediated mechanisms leading to therapeutic immune responses to cancer.

## Results

### Reconstructing intra-patient heterogeneity in single cell-derived organoid models

To study the generation of intra-tumor heterogeneity of CRC in PDTOs, we first engineered single cell-derived organoids that retained CRC characteristics (schematic, Fig. 1A). Freshly resected CRC tumor tissues were used to establish PDTOs as previously described(*19*). By immunohistochemistry, the patient tumor and matched PDTOs showed consistent staining for colorectal-specific nuclear markers, including caudal type homeobox 2 (CDX2) and cytokeratin 20 (CK20) as well as oncogenic drivers such as ERBB2 (Fig. 1B). Besides, we observed that PDTOs exhibit similar dMMR characteristics as the primary tumor by histological evaluation (Fig. 1B). These data suggested that the characteristics found in the patient tumor were retained in PDTOs.

**Fig. 1.**
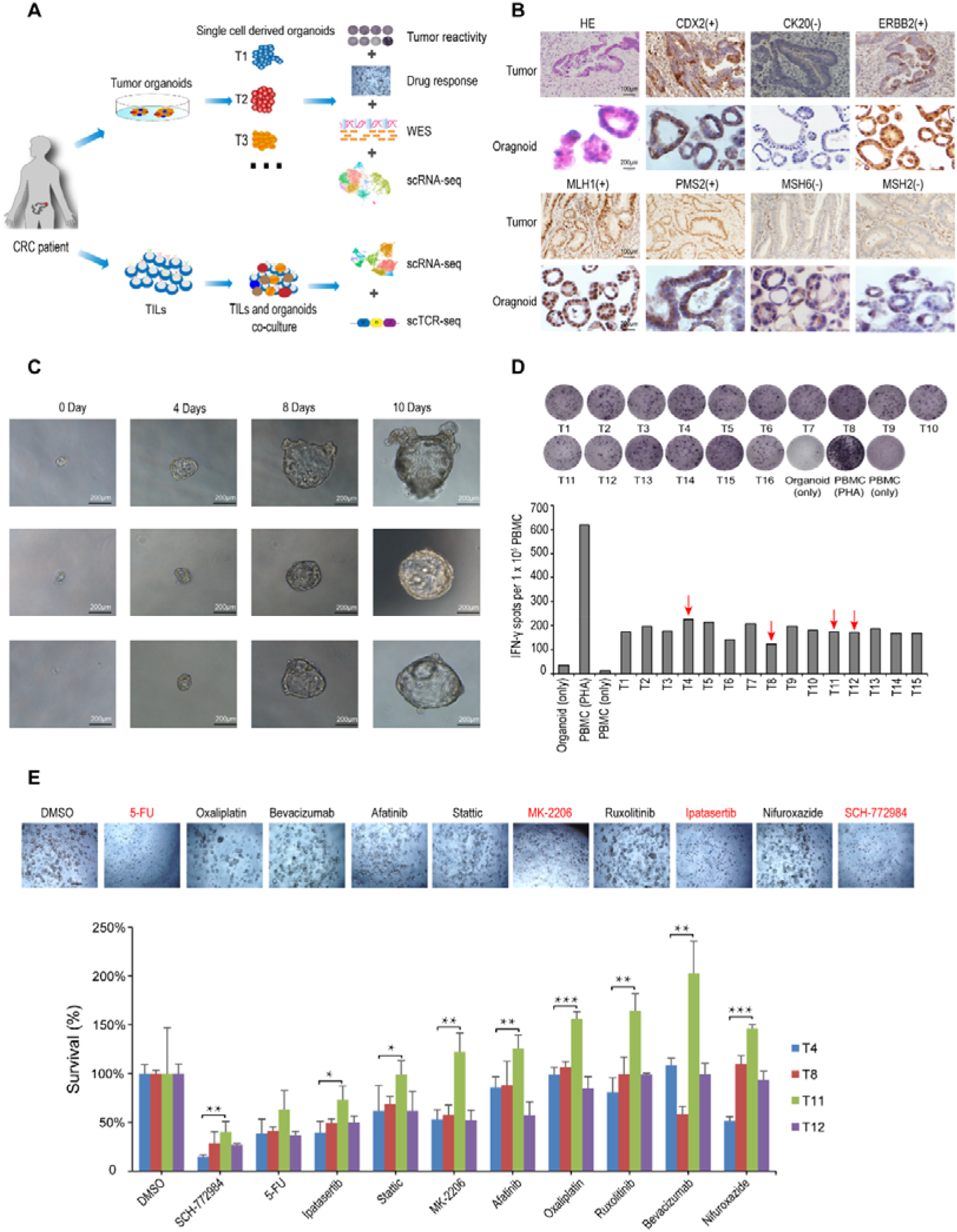
Generation of single cell-derived organoid and induction of tumor reactivity in PBMCs. (**A)** Schematic representation of the experimental strategy. (**B)** Representative images of H&E, and IHC of CDX2, CK20, ERBB2, MLH1, PMS2, MSH6, and MSH2 from primary tissue and patient-derived organoids. Scale bar = 100 μM in primary tissue images and 200 μM in organoid images. **(C)** Representative images of single cell-derived organoid morphology from 0, 4, 8, 10 days. **(D)** Quantification of organoid-induced IFN-γ production of PBMCs by an IFN-γ ELISPOT assay. Data are representative of a single experiment. **(E)** Microphotograph images of T11 organoids at 3 days treated with indicated drugs (up) and mean survival is displayed for exposure to indicated drugs (down) (n = 3). *P < 0.05, **P < 0.01 and ***P < 0.001 as determined by a student’s t-test.

Following the successful generation and characterization of PDTOs, their dissociation into single tumor cells further allowed the analyses of clonal growth, where each single cell-derived organoid clone showed a different growth rate and noticeable morphological differences (Fig. 1C). In total, 16 of 48 single cell-derived organoids were finally established, expanded, and cryopreserved for further experiments.

We next studied whether single cell-derived CRC organoids induced heterogeneous T cell tumor-reactive responses. Recently, Dijkstra et al. have reported that tumor organoids can be used to stimulate and propagate tumor-active T cells from PBMCs of CRC patients(*20*). Here, we established T cell co-culture assays with each single-cell-derived organoid clone and evaluated the tumor reactivity of T cells. Before co-culturing each clone with autologous PBMCs, they were pre-stimulated with IFN-γ to enhance antigen presentation. 16 out of 16 (100%) single cell-derived organoids, when co-cultured with autologous PBMCs, induced a significant PBMC IFN-γ secretion (≤ 12-fold increase) when compared to PBMCs alone (Fig. 1D). However, the magnitude of these responses varied among different single cell-derived CRC organoid clones, maybe due to intrinsic differences, including somatic mutations and antigen expression/presentation characteristics. Based on these IFN-γ tumor reactivity assay results, we defined and selected T4 (high reactivity), T8 (low reactivity), and T11 & T12 (medium reactivity) for further testing drug response.

To test the drug responses of organoid clones, chemotherapeutic and targeted therapy drugs including 5-Fluorouracil (5-FU), oxaliplatin, bevacizumab, nifuroxazide, SCH-772984 (ERK inhibitor), ipatasertib (AKT inhibitor), MK-2206 (AKT inhibitor), stattic (STAT Inhibitor), afatinib (ERBB2 target), and ruxolitinib (JAK inhibitor) were used to treat the selected organoid clones (T4, T8, T11, T12). The results showed that all the four organoid clones were sensitive to the SCH772984, 5-FU, MK-2206, and stattic. Besides, we found that the T11 organoid clone exhibited more resistance to targeted therapies than T4, T8, and T12 when organoids were treated with the same drug concentration (Fig. 1E). In addition, T4 organoid clones showed a lower survival ratio than other clones when treated with nifuroxazide, which can selectively mediate cell apoptosis in ALDH1-high melanoma cells(*21*). The cause of diversification of drug responses is still unclear, but a study showed no significant correlation with the degree of mutations(*5*). To better understand the CRC heterogeneity and diversification of drug responses, the four CRC organoid clones were analyzed by WES, RNA-seq, and scRNA-seq approaches.

### Genetic diversity is preserved between different organoid clones

Prior reports indicate that primary tumors and organoid cultures share 90% of somatic mutations(*19*). Interestingly, our somatic mutation analysis of tumor tissues, bulk organoids, and single cell-derived CRC organoids revealed commonality and heterogeneity in tumor somatic mutations and neoantigens. The results showed that a total of 2155 (71.74%) exonic mutations (SNVs and Indels) were shared among the bulk organoids and single cell-derived organoid clones (Fig. 2A), while unique mutations were also identified. The T4 organoid clone harbored significantly more unique mutations (270) than T8 (146), T11 (110), and T12 (118) (Fig. 2A). Based on SNVs and Indels, tumor neoantigens were predicted via NetMHCpan-3.0. Similarly, there were 246 predicted neoantigens shared among different organoid clones, while unique neoantigens could still be found in T4 (20), T8 (19), T11 (6), and T12 (27) (Fig. 2C). These results suggested that the diversity between organoid clones may also reside in their tumor-specific neoantigen presentation properties. Since single cell-derived CRC organoids may increase their genomic instability through passages, we analyzed the extent of somatic mutations in low and high passage numbers. Compared with passage1, passage 10 and 20 organoids acquire more than 119 specific somatic mutations (Fig. 2B), which may be due to the decreased ability to repair DNA during cell division in dMMR patients.

**Fig. 2.**
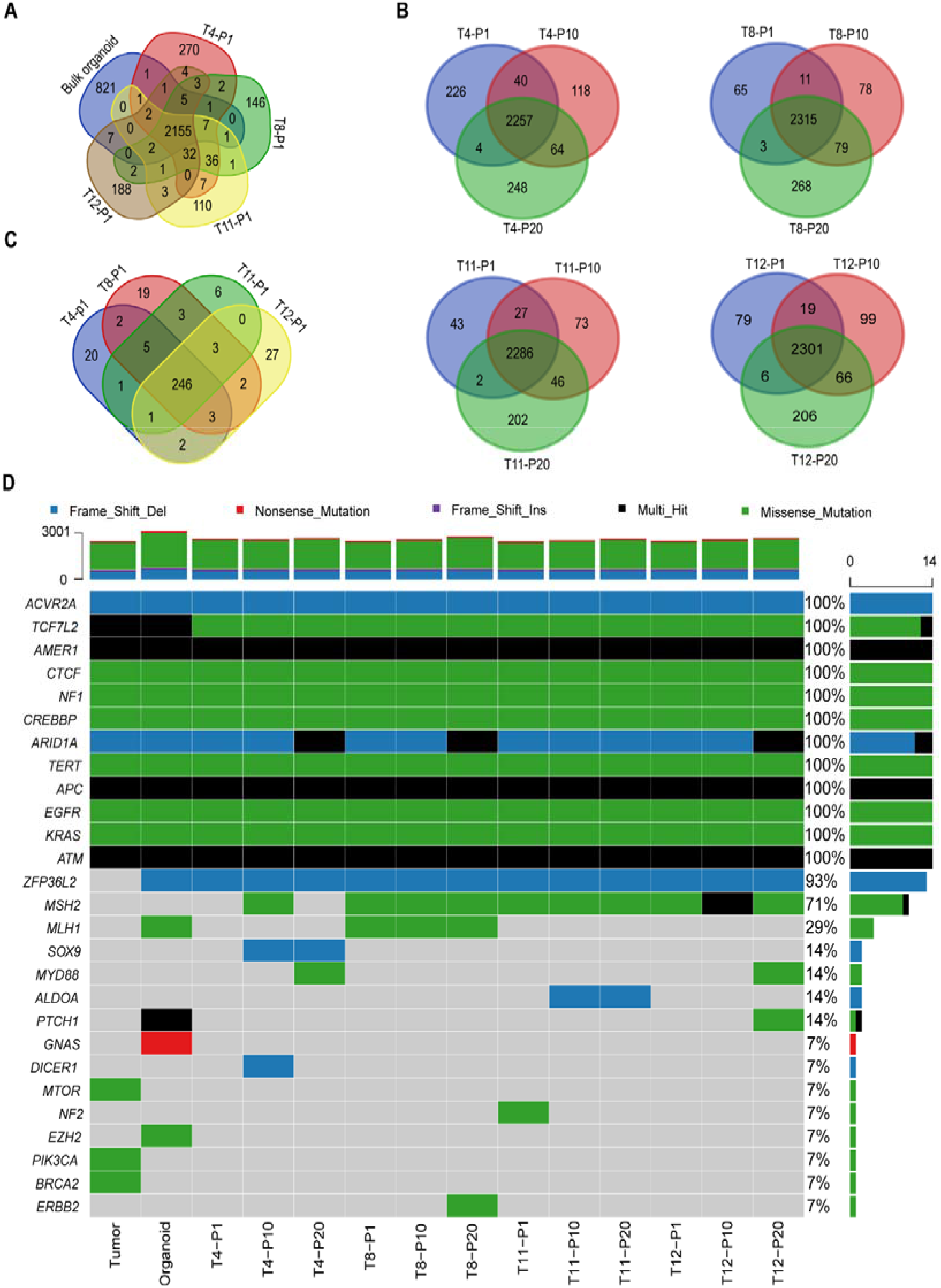
Genomic characterization of single-cell derived CRC organoids. **(A)** Venn diagram displaying the number of shared and unique mutations (SBS and INDELs) of bulk and clonal (T4, T8, T11, and T12) organoids. **(B)** Venn diagrams displaying the number of shared and unique mutations (SBS and INDELs) of organoid lines with different passages (P1, P10, and P20). **(C)** Venn diagrams displaying the predicted neoantigens of each CRC organoid line using the NetMHC-3.0 pan algorithm. **(D)** Oncoplot of the CRC driver gene mutations in tumor tissues, bulk organoids, and clonal organoids by WES sequencing.

Since tumor driver gene mutations could affect tumor growth or progression, we further examined the presence of driver gene mutations in the tumor tissue, bulk organoids, and single cell-derived organoids. In all cases, mutations were present for most of the CRC driver gene mutations (e.g., *APC, KRAS, EGFR, ATM, ARID1A*) analyzed using the COSMIC database (Fig. 2D). However, some other CRC driver gene mutations such as *MTOR, BRCA2*, and *PIK3CA* were only detected in the primary tumors, while *ZFP36L2, MLH1, PTCH1, GNAS*, and *EZH2* only existed in bulk or single cell-derived organoids, and this may be due to the pure cellularity achieved in organoid cultures (Fig. 2D). Interestingly, mutations in the cancer driver gene *MLH1* were specifically found in the organoid clone T8, but not in T4, T11, or T12 (Fig. 2D). In addition, other driver genes, including *ERBB2, SOX9, ALDOA, PTCH1, MYD88*, and *DICER1*, harbored unique mutations just found in CRC organoid clones passage 10 or 20, but not in passage 1. Taken together, these data indicated that organoids reconstructed tremendous intratumor heterogeneity and revealed that single cell-derived CRC organoids were not genetically identical.

### Single-cell transcriptome analysis reveals a high degree of cancer stem cell heterogeneity in different organoid clones

To further address the degree of organoid heterogeneity, we performed single-cell RNA sequencing of bulk tumor organoids and T4, T8, T11, T12 organoid clones (fig. S3A and 3B; table S1). The results obtained from the integrated data set revealed the existence of a total of nine cell subtypes: six stem cell subpopulations, two transient-amplifying cells (TACs) subpopulations, and one subpopulation of epithelial-mesenchymal transition (EMT)-like cells. These results were annotated based on cell-type-specific signatures and marker genes (Fig. 3C). Stem cell transcriptomes were defined by the specific expression of colorectal CSC markers such as *CD44, ITGB1*, and *SOX4*. TACs transcriptomes were assigned by the high expression of stemness genes and proliferation markers, including *MKI67* and *TOP2A* (Fig. 3D). EMT-like cell transcriptomes were defined by the high expression of *S100A4* and the low expression of *CDH1*.

**Fig. 3.**
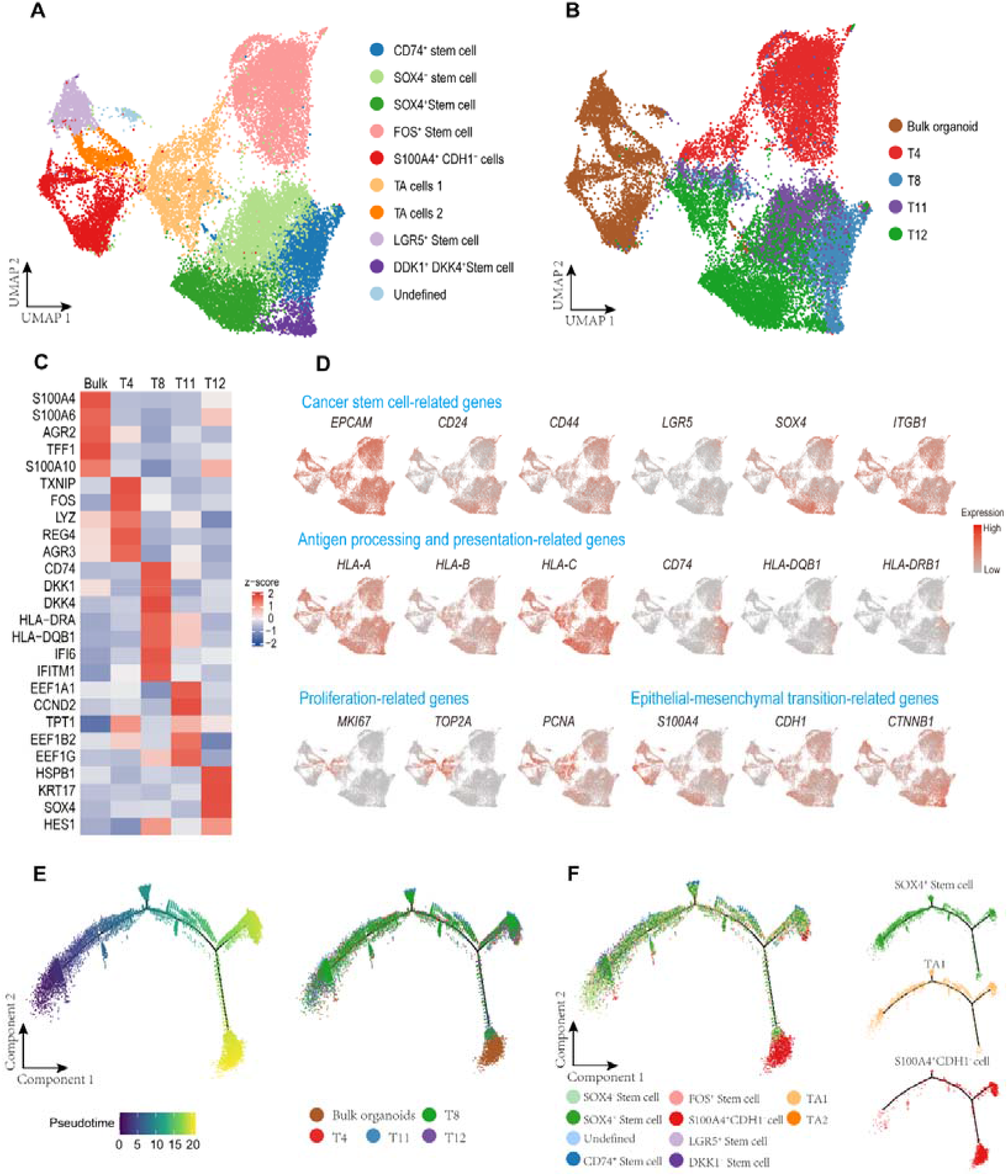
ScRNA-seq profiling of single-cell derived clonal colorectal organoids. **(A)** UMAPs of 24690 cells from bulk and clonal (T4, T8, T11, and T12) organoids. Cells are annotated into nine clusters based on transcriptome profiles. **(B)** UMAPs embedding all sequenced organoid cells, color-coded for organoids of origin. **(C)** Gene expression heat map of specific markers in bulk and clonal (T4, T8, T11, and T12) organoids. **(D)** Single-cell gene activities of cancer stem cell-related genes, antigen processing, presentation-related genes, and epithelial-mesenchymal transition-related genes. **(E)** Pseudotime analysis of bulk and clonal (T4, T8, T11, and T12) organoids, color-coded for organoids of origin. **(F)** Pseudotime analysis of organoids, cell subtypes are labeled by colors.

Bulk and single-cell derived organoids were mainly composed of epithelial cell types, enriched in *EPCAM* and *CDH1* markers, while they did not express fibroblast lineage-specific markers such as *VIM, COLLA1, COLLA2, LUM, FBLN1*, and *FBLN2*(*22*). The high expression of intestinal lineage markers, such as *MUC3, MUC5, MUC6, CDX1*, indicated that all cells were of strict intestinal origin.

The clones obtained from single cell-derived CRC organoids (T4, T8, T11, and T12) were composed of stem cells and TACs, with an absence of epithelial-mesenchymal transition (EMT)-like cells, suggesting that these organoid clones were in the state of stem cells or enterocyte progenitors (Fig.3A). Pseudotime analysis further showed that the stem cell clusters, mainly found in single cell-derived organoids, were only present at the beginning of the trajectory path. However, S100A4^+^ CDH1^-^ EMT-like cells found in the bulk organoids were only found at a terminal state (Fig.3F and fig. S1D). Therefore, compared to bulk organoids, single cell-derived organoids can better preserve stem cell phenotypes.

Concomitantly, single-cell transcriptome analysis revealed a significant degree of stem cell heterogeneity across the four organoid clones (T4, T8, T11, and T12). T4 cancer stem cells highly expressed the proto-oncogene *FOS* (Fig. 3C and table S3), which plays an essential role in tumor progression and promotes cancer stemness(*23*). The T11 clone highly expresses *eEF1A1*, which can inhibit P53, P73, and chemotherapy-induced apoptosis resulting in chemoresistance (Fig. 3C and fig. S1E). This increase may explain why the T11 clone showed increased resistance to drug treatment than the T4, T8, and T12 clones(*24*). Also, the higher expression of *ALDH1A1* in T4 may explain its increased sensitivity to nifuroxazide (Fig.S1E). T8 and T12 presented a cluster of *DDK1* and *DKK4* positive stem cells, absent in T4 and T11 organoid clones (Fig. 3C and fig. S1E). DKK proteins antagonize Wnt/ β-catenin signaling through binding to the LRP5/6 receptor. DKK1 has been involved in suppressing antitumor immune responses in CD8^+^ T cells through the GSK3β/E2F1/T-bet axis(*25*), suggesting that the identified DDK1^+^ DKK4^+^ CRC stem cell subpopulations may be associated with resistance to cancer immunotherapy. Moreover, the DEG analysis revealed an enrichment of genes involved in cell cycle-related pathways (e.g., DNA replication and Mismatch repair) in bulk organoids and the T4 clone. In contrast, the organoid clone T12 exhibited upregulation of genes belonging to colorectal cancer development-related pathways (e.g., WNT signaling pathway and PI3K−AKT signaling pathway) (fig. S1, A and B). These results showed that the single cell-derived organoids exhibit heterogenous stem cell-like characteristics, which might be ideal models for further studies on cancer stem cells.

Due to the relevance of major histocompatibility complex (MHC) class I and class II molecules in adaptive immune responses and the loss of heterozygosity (LOH) in MHC-I in up to 60% of dMMR CRC(*26*), we next studied their expression in our tumor organoids. Our single-cell transcriptome analysis demonstrated that MHC class I antigen processing and presentation-related genes such as *HLA-A, HLA-B*, and *HLA-C* were highly expressed in bulk and single cell-derived organoids (Fig. 3D). Furthermore, T4 and T8 organoid clones exhibited a high expression of markers associated with MHC class II antigen processing and presentation, such as *CD74, HLA-DRA*, and *HLA-DQB1*, suggesting that they may be more sensitive to MHC class II-mediated antitumor immunity by CD4^+^ T cells (Fig. 3D).

We also examined the protein expression of MHC class I and MHC class II molecules in organoids with or without interferon-gamma (IFN-γ) stimulation by western blotting. The results showed that bulk organoids and T4, T8, T11, and T12 organoid clones expressed both MHC I and MHC II proteins, and their expression level was significantly upregulated after IFN-γ stimulation (fig. S1C), consistent with prior studies(*20*). Overall, these data indicated the suitability of the tumor organoids established in our study for further investigating antitumor activities elicited by autologous T cells isolated from patients.

### scRNA-seq and scTCR-seq analysis reveals tumor-reactive CD4^+^ T cells are highly enriched in proliferating T cells

To enrich tumor-reactive T cells from tumor-infiltrating lymphocytes (TILs), we first expanded patients-derived TILs in vitro and then evaluated their antitumor activities towards the organoids using live imaging. The results showed that TILs generated substantial killing activities toward tumor bulk organoids (Extended Data Fig. 2e). To characterize the tumor-reactive T cells transcriptomic features and functional properties, we performed scRNA-seq and single-cell T-cell receptor (scTCR)-seq of TILs before and after co-culturing them with bulk tumor organoids. In total, we obtained scRNA-seq profiles from 10929 T cells, and 9748 (89%) paired TCR sequences (fig. S2A). Four CD4^+^ T clusters were identified based on scRNA-seq profiles, including Th1/2 cells, FOXP3^+^ regulatory T cells (T_reg_), effector memory T cells (T_EM_), and proliferating T cells which also had an exhausted phenotype based on exhaustion markers, such as *PDCD1, CTLA4*, and *LAG3* (Fig.4A and fig. S2C; table S2).

**Fig. 4.**
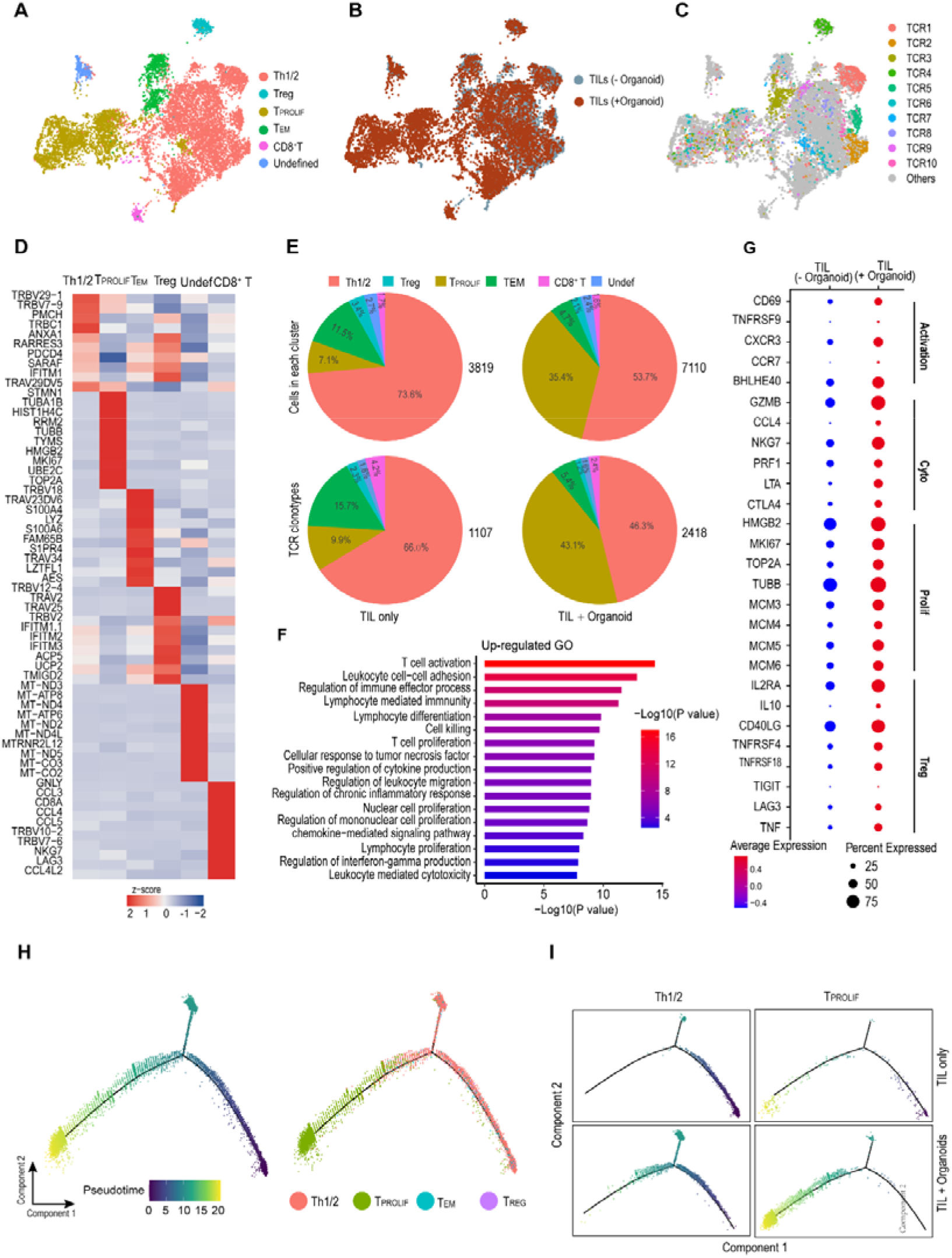
Single-cell RNA-seq analysis of TILs before and after co-culture with organoids. **(A)** UMAPs of all tumor-infiltrating T cells before and after co-culture with autologous tumor organoids, color-coded for cell types. **(B)** UMAP of tumor-infiltrating T cells colored by organoids treatment status. **(C)** UMAP of tumor-infiltrating T cells colored by TOP10 TCRαβ clonotype frequency. **(D)** Gene expression on UMAP embedding of specific markers that were used for cluster annotation. **(E)** Pie charts summarize the distribution changes of cell subsets (top) and TCR clonotypes (cell ≥ 3 cells) (down) before and after co-culture with autologous tumor organoids. **(F)**. Gene Ontology (GO) enrichment analysis of the upregulated genes (P<0.05) in TILs after co-culture with tumor organoids. **(G)** Bubble chart showing the expression levels and sizes of activation, cytotoxicity, proliferation, and regulatory T cells-related markers in TILs before and after co-culture with autologous tumor organoids. **(H)** Pseudotime analysis of CD4^+^ T cells from TILs before and after co-culture with tumor organoids. T cell subtypes are labeled by colors. **(I)** Pseudotime analysis of Th1/2 and T_PROLIF_ cells, from TILs (up) and organoids co-cultured TILs (down).

To study gene expression patterns associated with tumor-reactive CD4^+^ T cells, we first investigated the ratio of cell and gene expression changes in TILs after co-culture with tumor bulk organoids. Notably, the proportion of proliferating CD4+ T cells increased significantly from 7.1% to 35.4% after co-culture with organoids, suggesting that tumor-reactive T cells in the proliferating cluster clonally expanded upon recognition of tumor organoids (Fig.4, B and E). Pseudotime analysis also revealed the significant trajectory differences of TH1/2 and proliferating CD4^+^ T cells in organoid treated TILs (Fig.4, H and I). T cell activation related genes (*CD69, TNFRSF9, CXCR3, CCR7*, and *BHLHE40*), cytotoxicity related genes (*GZMA, GZMB, CCL4, NKG7, PRF1, LTA, GNLY*, and *CTLA4*), proliferation-related genes (*HMGB2, MKI67, TOP2A, TUBB, MCM3, MCM4, MCM5*, and *MCM6*), T cell recruitment chemokines (*CCL3, CCL4*, and *CXCL13*) and regulatory T cell-related genes (*IL2RA, IL-10, CD40LG, TNFRSF4, TNFRSF18, TIGIT, LAG3*, and *TNF*) were significantly upregulated (p-value < 0.01, Wilcoxon test, and fold change ≥2) (Fig. 4G). We also identified CD4^+^ BHLHE40^+^ TH1 cells, which feature MSI tumors and are more sensitive to ICB therapy(*27*). GO and KEGG enrichment analysis of DEGs showed enrichment of genes in CD4^+^ T cells involved in T cell activation, cytotoxicity, and T cell proliferation-related pathways, indicating that upon co-culture with tumor bulk organoids, TILs become fully activated (Fig. 4F and fig. S2B; table S4).

We next assessed changes in the potential specificity of TILs to tumor antigens by studying if their TCR clonotypes and abundance changed after co-culturing them with tumor organoids. Using correlative analysis of scRNA-seq and scTCR-seq data, we found that the top frequent TCR clonotypes tend to distribute among clusters (Fig. 4C), in accordance with previous studies(*27-29*). Tumor antigen-specific CD4^+^ T cells substantially expand clonally during immune responses, and proliferating CD4^+^ T cells (MKI67^+^) cells are predictive response markers to checkpoint blockade immunotherapies in thymic epithelial tumors and non-small cell lung cancer(*14, 30*). Our results showed that after co-culturing TILs with tumor organoids, their TCR clonotypes (clonotype defined as ≥3 cells sharing the same TCRα/β pair) in the proliferating CD4 T cell cluster significantly increased from 110 to 1043. We also noted a significant increase in the CD4 clonal abundance of TCR2, TCR3, TCR4, TCR6, TCR6, and TCR15 in T_PROLIF_, while many novel clonotypes emerged, expanding after co-culture, suggesting that these clonotypes may be reactive to tumor organoids (Fig. 5A). These data were consistent with another study on the characterization of the CD4^+^ cytotoxic T lymphocytes by scRNA-seq(*31*). In contrast, no significant changes in TCR clonotype and frequency were observed in Th1/2, T_EM_, and Treg (Fig.4, C and E). Overall, our results provided evidence that tumor-active T cells may be enriched in the proliferating CD4^+^ T cells.

**Fig. 5.**
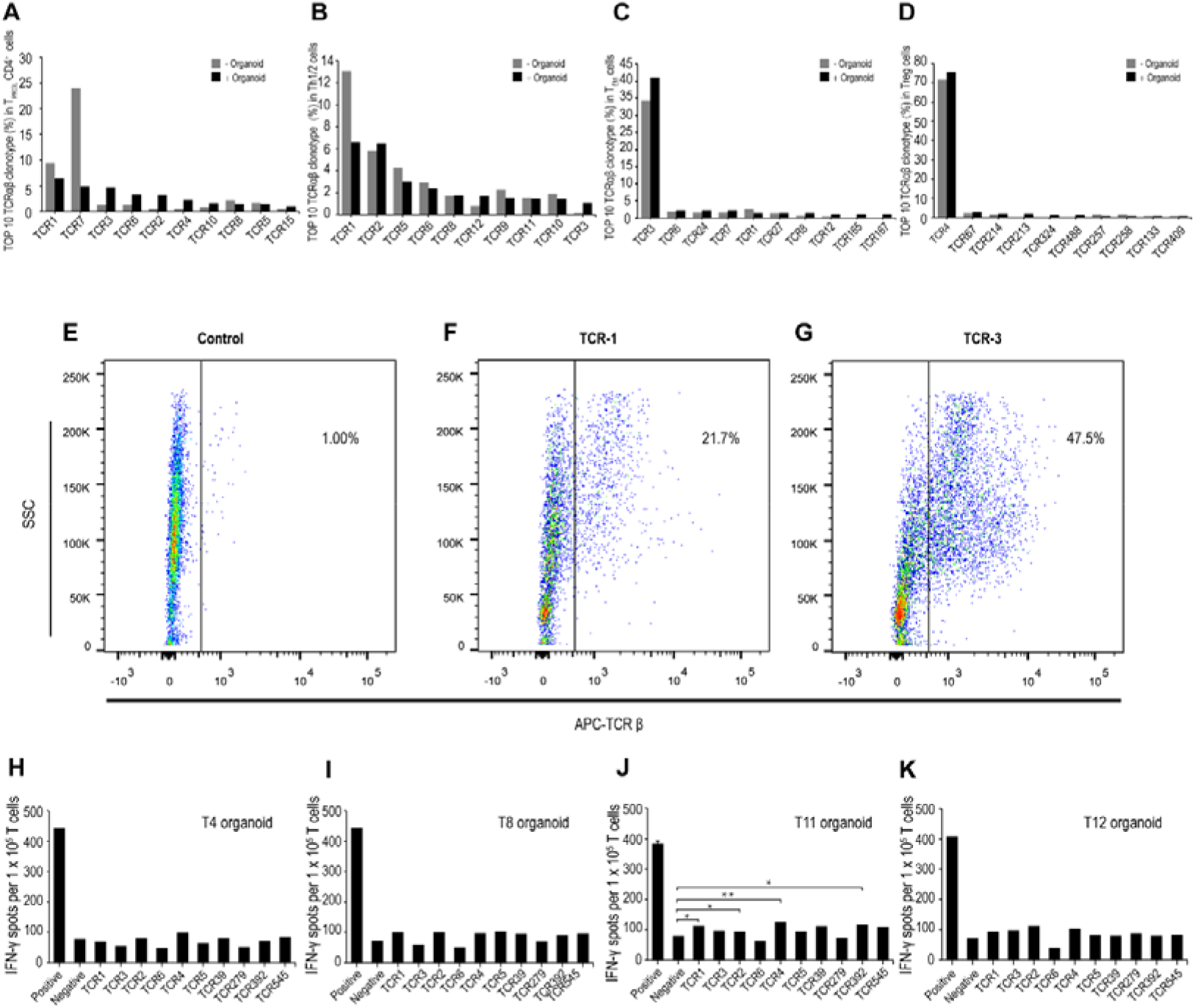
Identification of tumor-active CD4^+^ TCRs in CRC. **(A)** Bar chart showing the top10 TCRαβ clonotype frequency in proliferating CD4 T cells colored by organoids treatment status. **(B)** Bar chart showing the top10 TCRαβ clonotype frequency in proliferating Th1/2 cells colored by organoids treatment status. **(C)** Bar chart showing the top10 TCRαβ clonotype frequency in T_EM_ cells colored by organoids treatment status. **(D)** Bar chart showing the top10 TCRαβ clonotype frequency in Treg cells colored by organoids treatment status. **(E to G)** Representative β gene expression in primary T cells was determined by flow cytometry. T cells without **exogenous TCR** transfection were used as control. **(H to K)** Numbers of spot-forming cells per 1.5 × 10^5^ TCR electroporated T cells co-culture with T4, T8, T11, T12 organoid clones using an IFN-γ ELISPOT assay. Data are represented as mean (n = 2). *P < 0.05, **P < 0.01 and ***P < 0.001 as determined by a student’s t-test.

To define the potential autologous tumor-antigen recognition of the top ten TCR clonotypes identified in the CD4 T cells co-cultured with tumor organoids, we cloned those TCR sequences into expression vectors followed by *in-vitro* transcription. T cells from healthy HLA-A*11:01 donors were individually electroporated with each one of the ten selected TCR mRNAs. Their exogenous expression was determined after 24 h, which showed highly efficient transfection of 20% to 50 % in primary T cells (Fig. 5D to F). We next assessed the levels of IFN-γrelease as a predictive value of the anti-tumor response of T cells expressing the different TCR clonotypes. To this end, TCR-engineered T cells were co-cultured with four different single-cell derived organoid clones, and IFN-γlevels secreted by CD4 T cells were measured by ELISPOT assays. TCR-engineered T cells displayed divergent reactivities to single-cell organoids. For example, TCR1, TCR2, TCR4, and TCR392 exhibited stronger reactivities against T11 than T4, T8, and T12 organoid clones (Fig. 5, H to K). In total, 4 out of 10 TCRs exhibited tumor recognition towards the organoids (Fig. 5J), and 3 of the 4 TCRs showed significantly increased clonal abundance after co-culture with tumor organoids.

The use of tumor organoid-TILs co-culture assays efficiently leads to the clonal expansion of proliferating CD4+ T cells containing tumor-reactive T cell clonotypes. Overall, these results suggest that the combined analysis of scRNA-seq and scTCR-seq could facilitate the identification of potential tumor-reactive T cells. These platforms might provide relevant information for identifying TCR sequences with profound implications for developing personalized TCR-engineered cell therapies against tumors.

## Discussion

Recently, the systematic and integrated profiling of single cell-derived organoid clones’ genetic, transcriptomic, and functional responses became a better strategy to study tumor heterogeneity than individual cells, enabling the comprehensive description of almost all somatic mutations present in multiple single cells(*5*). In this study, we generated single cell-derived organoids in CRC to study their heterogeneity and tumor reactivity at the single-cell level along with their genomic characterization. We also typified different cell population types the cell types and differentiation states using droplet-based single-cell sequencing of transcriptomes. Moreover, by coupling the organoid-TILs co-culture model, high-resolution single-cell profiling of CD4^+^ TILs, and specificity recognition testing of candidate TCRs, we determined the properties associated with the cytotoxic CD4^+^ T cells.

Many studies have revealed that the original genetic diversity of the tumor was preserved in organoid cultures(*8, 19*). However, our results showed that while some driver gene mutations such as *PIK3CA, BRCA2*, and *MTOR* are observed in tumor biopsies, they are not present in bulk or single cell-derived organoids. Other driver gene mutations, including *ALDOA, MSH2, MLH1, ALDOA*, and *MYD88*, were only detected in organoid clones. In addition, it seems that a higher allele frequency of driver gene mutations exists in passage 10 and 20 organoid clones than in passage 1 organoids. This difference may reside in the enrichment or depletion of sub-clonal populations in organoids during their cell culture passages and the potential acquisition of new mutations due to defects in mismatch repair processes. In any event, the unique somatic mutations that exist in each organoid clone finally contribute to the diversity of neo-epitopes, in agreement with the differences found in HLA class I peptide presentation between organoid clones in a microsatellite stable (MSS) CRC patient (*11*). These results revealed that the organoid genomes might not be stable with increasing culturing passages, especially in MSI CRC patients, and it may be more reasonable to use early-passage (<10) tumor organoids for experiments.

Previous studies in microsatellite instability (MSI) CRC patients showed that the single cell-derived organoids with particular somatic mutations displayed marked differences in drug responses, even some closely related organoid clones also had substantial and reproducible differences(*5*). Because single cell-derived organoids are derived from cancer stem cells in CRC, this provides a unique perspective that the stem cell heterogeneity may also underly the diversification of drug responses(*32*). Our studies revealed a unique stem cell subtype with high expression of DDK1 and DKK4 in both T8 and T11 organoid clones, which may be associated with resistance to cancer immunotherapy. Other stem cell subtypes in T8 organoids highly express *CD74, HLA-DRA*, and *HLA-DQB1*, which are related to the major histocompatibility complex (MHC) class II antigen processing and presentation, may be more sensitivity to CD4-mediated anti-tumor immune response.

The mutation-specific CD4^+^ cytotoxic T lymphocytes (CD4-CTLs) have been reported to mediate anti-tumor cytotoxicity in human bladder cancer and epithelial cancer(*14, 15*). The autologous T cells and tumor organoids co-culture platform has demonstrated the ability of unbiased and systematic analysis of T cell-mediated tumor recognition(*20*). In our study, we used an autologous TILs and tumor organoids co-culture platform, as well as scRNA-seq and scTCR-seq, to study the relationship between phenotypic characteristics and TCR with anti-tumor reactivity properties in cytotoxic CD4^+^ T cells. Furthermore, there are already some single-cell transcriptome analyses to characterize cytotoxic T cells. CD4-CTLs were reported to express a high level of cytotoxicity-related genes such as *CX3CR1, GPR56, CD244, CD314, KLRG1, GZMB*, and *PRF1*, and enriched in the CD4-T_EMRA_ subset in donors with previous DENV and CMV infection(*31*). Another study showed that multiple cytotoxic CD4+ T cell states are enriched and clonally expanded in bladder tumors and possess lytic capacity against tumors(*14*). Based on those previous studies, we considered that CD4 cytotoxic T cells may have the characteristics of T_EMRA_ phenotype and clonal expansion and screened for potential cytotoxic CD4^+^ T cells. In line with recent studies, we observed that 55.87% TILs express CD4^+^ T_EMRA_ phenotype (defined as CD3^+^CD4^+^CD45RA^+^CCR7^−^ cells) and a high level of cytotoxic related genes *GZMA, GZMB, NKG7* and *PRF* (fig. S2C). Furthermore, our results showed that the proliferating CD4^+^ T cell subsets were obviously clonal expanded after co-culture with autologous organoids, suggesting that these restricted TCR repertoires may be the result of recognition of MHC class-II antigens which may include autologous tumor antigens.

It has been reported that CTLA4 blockade can rapidly increase the number of activated effector CD4^+^ T cells and CD4^+^ FoxP3^+^ Tregs, revealing that the proliferating CD4^+^ T cells may contain both cytotoxic cells and regulatory cells(*29, 31*). However, we did not detect CD4^+^FoxP3^+^ Tregs in the proliferating CD4^+^ T cells population. Besides, the proportion of non-proliferating Tregs in TILs before and after co-culture with organoids was 3.4% and 2.1%, respectively, which showed no significant change (Fig. 4E).

To further analyze the tumor activity of those restricted TCRs with clonal expansion after coculture with tumor organoids in proliferative T cells, we used recognition testing against patient-derived organoids to define the tumor reactivity of CD4^+^ TCR clonotypes infiltrating the tumor. We found that 4 of 10 (40%) TCR exhibited tumor recognition (Fig. 5J). Finally, we identified CD4 cytotoxic T cells as a group of proliferative CD4^+^ T cells with T_EMRA_ phenotype and clonal expansion after co-cultured with autologous tumor organoids. A recent study showed that 102 of 123 (83%) CD8^+^ Tex-TCRs analyzed across 4 metastatic melanoma patients were confirmed to be tumor-specific and active(*33*). However, when the TCR-T cells were cocultured with organoids, we did not observe the obvious death of tumor cells. The possible reason for this discrepancy may reside in the low expression of the tumor-specific antigens or insufficient secretion of relevant cytotoxic cytokines. The co-culture of organoids with a single immune cell type may not evaluate the complex interaction between immune cells and tumor cells, and new platforms need to be designed.

In summary, our data indicate that the single cell-derived organoids are mainly composed of cancer stem cells and TACs and are highly heterogeneous in terms of somatic mutations, driver genes, and stem cell subpopulations. Single cell-derived organoids may be an ideal model for studying immunotherapy that targets cancer stem cells. The antitumor recognition mediated by CD4-CTLs can be effectively identified and screened from the proliferating CD4^+^ T cells with T_EMRA_ phenotype and clonal expansion using autologous T cells and tumor organoids co-culture model. A combination of transfer of gene-modified T cells armed with CD4 T_PROLIF_-TCRs cells and chemotherapeutic targeting specific mutation clones could result in effective, durable, and personalized tumor cytotoxicity.

## Methods

### Human subjects

The study was approved by the Ethical Committee of the BGI-Shenzhen - Sun Yat-Sen University Cancer Center. The PBMC, organoid lines, and TILs used in this study were derived from a female CRC patient (provided by Sun Yat-Sen University Cancer Center, Guangzhou, China). Mismatch repair deficiency was performed by immunohistochemical staining for MSH2, MSH6, MLH1, and PMS2, while microsatellite instability (MSI) analysis was confirmed based on PCR amplification of the 5 microsatellite loci BAT-25, BAT-26, D5S346, D2S123, and D17S250.

### CRC organoid culture

Surgically resected CRC tissue or biopsy tissue was obtained and processed within 24 hours. Patient-derived CRC organoids were established as previously described(*19*). Briefly, tumor tissue was washed with ice-cold phosphate-buffered saline (PBS) buffer and minced into small tumor pieces. The fragments were digested using 125 μg/ml dispase type II (Invitrogen) and 1.5 mg/mL collagenase II at 37°C for 1 hour. Tumor cells were filtered with a 100-μm cell strainer and centrifuged at 350g for 5 min before embedding Matrigel. After incubation for 15-30 min at 37°C, tumor organoids were cultured in Advanced DMEM/F12 supplemented with 1x N2 (Gibco), 1x B27, 500 ng/ml R-Spondin-1, 100 ng/ml Wnt3a, 100 ng/ml Noggin, 10 nM Gastrin I, 50 ng/ml human recombinant EGF, 5 μM SB202190, 0.5 μM A83-01, 10 ng/ml human recombinant FGF-10, 10 mM Nicotinamide, 1 mM N-acetylcysteine, 2 mM GlutaMAX, 10 mM HEPES, 1 X Pen/Strep and 10 mM Y-27632 (Supplementary Table S5). In subsequent passages and expansions, CRC organoids were cultured in a medium without Wnt-3A.

### TILs generation

Tumor tissue was cut into 1-3 mm^3^ fragments and put in a digestion medium consisting of 10 mg/ml collagenase I and 10 mg/ml hyaluronidase at 37°C for about 40 minutes. TILs were passed through100-μm cell strainers and then centrifuged at 350g for 5 min. Subsequently, TILs were placed in 6 well plates with 2 mL of TIL media (RPMI 1640) supplemented with 10% FBS, 2 mM L–glutamine, 10 mM HEPES, 1 x Pen/Strep (Gibco), 50mM β-mercaptoethanol, and 6000 IU/ml human recombinant IL-2. Half of the media was removed and replaced every 2 to 3 days and cultures were passaged by adding fresh TIL media to maintain a cell density of 2 × 10^6^ cells/ml.

### Generation of single cell-derived organoid

CRC organoids were passaged and cultured for 1-2 weeks, 10-20 individual organoids were collected and dissociated by TrypLE Express (Thermo Fisher), centrifuged at 300g for 5 min and washed twice with PBS. Single cells were suspended in 1 mL DMEM-F12 media (no Wnt3A), and counted with an automated cell counter, adjusting cell density to 5 × 10^3^ cells/ml. 10 μL cell suspension volume was aspirated and embedded with 1 mL Matrigel. Clonal tumor organoid was cultured in 48-well plates at 20 μL per well. Changing the medium once a week and cells were observed through a light microscope every 2-3 days until the clonal organoids formed. Single clonal organoids were picked out, expanded, and cryopreserved for further studies.

### Evaluation of single-cell organoids reactivity

Co-culture of single cell-derived organoids and autologous PBMC were performed to evaluate the tumor reactivity. Briefly, one day before co-culture, PBMCs were suspended at a density of 2*10^^6^/ml in T cell medium (RPMI 1640 medium supplemented with 10% FBS, 25 mM HEPES, 2 mM l-glutamine, and 300 IU/ml IL-2) and cultured overnight at 37 °C with 5% CO2. Single cell-derived organoids were stimulated with 200ng/mL human recombinant IFN-γ for 24 hours before co-culture. The 96 well plates used for IFN-γ enzyme-linked immunospot (ELISPOT) assay were first washed 4 times with sterile PBS, then coated with 5 mg/mL anti-CD28 antibody, and kept at 4°C overnight.

The next day, each organoid clone was digested into single cells by TrypLE Express and resuspended in a T cell medium. The ELISPOT plate coated with anti-CD28 was washed twice with sterile PBS and blocked with 10% FBS for another 30 minutes at room temperature. PBMC were seeded at a density of 1*10^5^ cells/well and cocultured with tumor organoids for 24 hours at a 2:1 effector: target ratio. In addition, PBMC or organoid alone was used for negative control. PBMC cultured with anti-CD3 antibody (clone: OKT3, 10 μ g/ml) was used for positive control. IFN-γ ELISPOT assay was performed according to the manufacturer’s protocol. Spots were imaged and counted using an ELISPOT reader and associated software.

### Organoid CellTiter-Glo viability assay

The organoid clones (T4, T8, T11, and T12) were collected and dissociated into single cells by TrypLE Express (Thermo Fisher). The cell suspension was resuspended in 50% Matrigel50% complete medium. 1*104 cells each well were seeded in a 96-well plate and cultured for 24h. Drug screening was carried out using 5-FU (10μg/mL, Selleck), SCH772984 (5μM, Selleck), ipatasertib (5μM, Selleck), MK-2206 (3μM, Selleck), stattic (3μM, Selleck), afatinib (1μM, Selleck), oxaliplatin (2μM, MCE), ruxolitinib (0.3μM, Selleck), bevacizumab (3μM, MCE), and nifuroxazide (5μM, MCE). To test the drug response to organoid clones, cells were cultured for 3 days and cell viability was measured using CellTiter-Glo® 3D Cell viability assay (Promega) according to the manufacturer’s protocol. Survival ratios of drug-treated tumor organoids were evaluated by normalizing to the DMSO control (100%).

### Exome DNA sequencing and RNA sequencing

Genomic DNA and total RNA were isolated from normal tissue, tumor tissue, patient-derived CRC organoids, and single cell-derived organoids with different passages using the DNEasy kit (QIAGEN). For exome sequencing, DNA exome capture libraries were made by using the BGI Human All Exon V4 kit. Captured products were circularized and rolling circle amplification (RCA) was performed to generate DNA Nanoballs (DNBs). RNA-seq libraries were prepared using MGIEasy RNA Directional Library Prep Kit (MGI). The Exome sequencing and RNA sequencing performed pair-end 100bp sequencing on BGISEQ-500 or BGISEQ-T7 platform.

### Organoids-TIL co-culture

Autologous TILs were rested in RPMI 1640 medium supplemented 10% 10% FBS, 2 mM L–glutamine, 10 mM HEPES, 1 X Pen/Strep (Gibco) for one week. Tumor organoids were pretreated with 200ng/mL human recombinant IFN-γ for 24 hours before coculture. A total of 1*10^6^ TILs were cocultured with or without tumor organoids in a six-well plate for 24 hours at a 5:1 effector: target ratio. After 24 hours of coculture, TILs were purified using a human CD3 T cell isolation kit to perform scRNA and V(D)J sequencing.

### Single-cell RNA sequencing (scRNA-seq) and single-cell TCR-sequencing (scTCR-seq)

For scRNA and scTCR sequencing, the libraries of TILs before and after co-culture with tumor organoids were produced by a previously established protocol using the 10x Chromium Single-Cell Immune Profiling Solution assay. In brief, the TILs were resuspended in PBS with a final concentration of 100–800 cells/ μL. After cell barcoding and reverse transcription in droplets, cDNA was purified using Dynabeads MyOne SILANE and followed by PCR amplification. scRNA-seq libraries were generated using ChromiumTM Single-Cell 5’ Library & Gel Beads Kit and TCR enriched libraries were prepared using ChromiumTM Single-Cell V(D)J Enrichment Kit with amplified cDNA. The scRNA-seq and scTCR-seq libraries were sequenced on the BGISEQ-500 platform (MGI, Shenzhen). The scRNA-seq of bulk organoids and organoid clones was performed using the DNBelab C4 system as previously described(*34*).

### Transient expression of exogenous TCRs on primary T cells

Ten paired TCRα /β sequences were selected by tracking significantly expanded T cell receptor clones after being co-cultured with tumor organoids. The genes encoding TCR variable regions from single-cell sequencing were fused with murine TCR-α and -β2 constant regions respectively for reducing the effects of TCR mispairing. The TCR-α and -β chains were linked by a T2A self-cleaving peptide for optimal TCR expression as previously described(*35*). TCR α/β sequences were codon-optimized, synthesized (GenScript), and cloned into the T7-EGFP vector (Addgene). Plasmids encoding the TCR α and β genes were linearized with Spe I and then used for in vitro mRNA transcription using mMESSAGE mMACHINE T7 Transcription Kit (ThermoFisher Scientific) according to the manufacturer’s instructions.

Electroporation of primary T cells was conducted using the Celetrix electroporation system. Primary T cells were isolated from HLA-A1101 healthy donor PBMC using a human CD3 T cell isolation kit (Miltenyi) and resuspended in electroporation buffer at concentrations of ∼5×10^7^/ml. For each electroporation, ∼1×10^6^ T cells were mixed with 5 μg TCR α/β mRNA, aspirated into the cuvette and subjected to electroporation immediately, using the following electroporation conditions: 425 V, square-wave, 20-ms pulse width, and single pulse. The transfected cells were then transferred to pre-warmed HIPP™-T009 medium (Bioengine) supplemented with 2% FBS (Hyclone), 10 ng/mL IL-2, IL-7, and IL-15 (R&D systems), and cultured at 37 °C with 5% CO_2_ for 24 hours. To measure exogenous TCR expression, T cells transfected with TCR mRNA were washed, suspended in PBS containing 5% FBS at concentrations of ∼1×10^6^ cells/mL, and stained with APC-conjugated anti-mouse TCRβ chain antibody (Biolegend). Flow cytometry analysis was performed using BD FACSAriaII (BD), and data were analyzed using FlowJo10 (TreeStar).

To evaluate the antitumor reactivity of candidate TCRs, the TCR-engineered T cells were co-cultured with organoids and evaluated by IFN-γ ELISPOT method. Briefly, 1*10^^5^ TCR-transduced T cells were co-cultured with 1*10^5^ IFN-γ pretreated organoids in the ELISPOT plate. T cells without electroporation but coculture with organoids were used for negative control. Anti-CD3 antibody (clone: OKT3, 10 μg/ml) was added as positive control. IFN-γ ELISPOT assay was performed according to the manufacturer’s protocol.

### Data analysis

#### Whole exome sequencing and mutation identification

The WES library construction and sequencing methods were consistent with those previously described(*36*). The library was sequenced on the DNB-seq T7 platform with paired-end 100 bp mode. After base calling, the raw reads of each sample were filtered using Fastp software(*37*) and mapped to the human hg19 reference genome with the Sentieon genomics tools(*38*). Strelka2(*39*) was used for the detection of somatic SNVs and Indels with paired tumor-normal samples. The mutations were then annotated with ANNOVAR(*40*), and the mutations were removed with the following filtrations: (1) The frequency in known databases (the Exome Aggregation Consortium, 1000Genomes) did not exceed 0.5%; (2) The mutation was not in the repetitive region of the genome (annotated by ANNOVAR(*41*)); (3) The mutations in intergenic and intron regions were removed.

#### Neoantigen prediction

Mutant peptides were generated from non-silent SNVs or Indels. Mutant peptides which have the same sequences as peptides from NCBI Reference Sequence Database were filtered out. The affinity of each mutant peptide and corresponding HLA alleles from the same patient was predicted by NetMHC(*42*), NetMHCpan(*43*), PickPocket(*44*), PSSMHCpan(*45*), and SMM(*46*). The peptide was retained if there are more than two software-predicted affinity values with IC50s < 500.

#### Single-cell RNA-seq data processing

Single-cell data were obtained from two platforms, including 10X and iDrop. Single-cell data were divided into 2 separate groups for subsequent analysis, including TIL (only) and TILs (organoids treatment) groups, and the organoid group (bulk, T4, T8, T11, and T12). For the data (TILs (only) and TILs (organoids treatment)) generated by the 10X platform, the sequencing data were mapped to the human reference genome (GRCh38) using the Cell Ranger (version 3.0.2) and generated the unique molecular identifier (UMI) matrix. On the other hand, the iDrop C4 platform sequencing data (organoids group) was mapped to the human genome (GRCh38) by using STAR(*47*). And the UMI matrix for each sample was generated by PISA (https://github.com/pisa-engine/pisa).

Single-cell analysis of these data was performed using Seurat (version 4.0.4)(*48, 49*). First, cells with less than 500 genes or UMI less than 1,000 or mitochondrial ratio greater than 10% were removed for TILs (only) and TILs (organoids treatment) samples, cells with abnormally low and abnormally high genes were removed. The value of abnormally low genes: Lower quartile (Q1) of genes - 1.5 * InterQuartile Range (IQR) of genes. The value of abnormally high genes: Upper quartile (Q3) of genes – 1.5 * InterQuartile Range (IQR) of genes. IQR: Q3 – Q1.). After quality control, a total of 35,619 cells remained, including 24,690 organoid cells and 3,819 TILs (only) cells, and 7,110 TILs + Organoids cells. Then 2,000 variable genes were selected using the “FindVariableGenes” function for each group. The reciprocal PCA (rPCA) was used to integrate the TILs (only) and TILs (organoids treatment) groups by using “FindIntegrationAnchors” and “IntegrateData” functions. Organoid groups do not perform this procedure, but instead, merge multiple samples using the “merge” function. The linear dimensional reduction was performed by using the “RunPCA” function and 30 PCs were selected. Cell clustering and Uniform Manifold Approximation and Projection (UMAP) visualization were performed using the “FindClusters” and “RunUMAP” functions, respectively. The “FindAllMarkers” function was used to determine the marker genes of each population. And identify cell types were identified by the known marker genes.

#### TCR analysis

The TCR sequences, including TILs (only) and TILs + Organoids, were mapped to the human reference genome (GRCh38), assembled, and annotated by using the Cell Ranger (version 3.0.2) VDJ pipeline. We then mapped the TCR annotation results to individual cells to generate TCRαβ clonotype information for each cell. The top 10 TCR sequences were screened by sorting according to the number of TCR. Then the candidate TCR sequences were generated by comparing the TCRαβ clonotype of TILs (only) and TILs + Organoids.

#### Differential expression and functional enrichment analysis

Differential expression analysis for each cluster or each sample was performed using the Wilcoxon rank-sum test as implemented in the “FindAllMarkers” or “FindMarkers” function of Seurat (version 4.0.4)(*49*). For each cluster or each sample, only genes that met these screening conditions were considered as differentially expressed genes (DEGs):

1. Log2 fold change > 0.25;
2. p < 0.05;
3. min.pct > 0.25 (expressed in at least 25% cells in either group of cells).

#### Gene Ontology (GO) analysis and KEGG pathway enrichment analysis

Gene Ontology (GO) analysis and KEGG pathway enrichment analysis were performed using the R software clusterProfiler for DEGs in each cluster or sample(*50*). The annotation Dbi R package org.Hs.eg.db was used to convert gene symbol to gene id. A P-value less than 0.05 was taken to indicate statistical significance.

#### Development trajectory analysis

Development trajectory analysis was performed on a subset cluster of TIL group cells (CD4^+^ T cells including Th1/2, T_PROLIF_, T_EM,_ and Treg) and all Organoid group cells (Bulk, T4, T8, T11, and T12) by using the Monocle2 R package(*51*). The variation in each gene’s expression across cells varies according to the mean was performed by using the “dispersionTable” function. And the cell’s progress related genes were defined by using “subset” function with parameter “mean_expression >= 1 & dispersion_empirical >= dispersion_fit”. Dimensionality reduction was performed using the “reduceDimension” with default parameters, using the cell’s progress-related genes. Finally, the “orderCells” function was used to order the cells and construct the developmental trajectory.

### Statistical analysis

For mutation visualization and statistics, we used maftools (v2.6.05) and ggplot2 (v3.3.3), and all analyses were performed in R studio (R 4.0.2). For single-cell RNA-seq, statistical analyses were performed using R (version 4.0.4). Graphs were generated using the R package (ggplot2 and ComplexHeatmap). The Wilcoxon rank-sum test was used to calculate differential genes between different cell types or samples, and p < 0.05 was considered significant. Fisher’s Exact Test was used for gene function annotation.

The phenotyping experiments of TIL by flow cytometry, HE staining, and immunohistochemistry were performed once. Drug response assays are representative of at least two independent experiments and were performed in three replicates. Tumor reactivity assay and TCR functional experiments were performed without duplicates due to material restrictions.

## Data accession

The data supporting the findings of this study have been deposited in the CNSA (https://db.cngb.org/cnsa/) of CNGBdb with accession number CNP0002111. All data analyzed during this study have been included in this published article and its supplementary information files.

## Author Contributions

Conceptualization: JY, MPM, and XZ. Methodology and Investigation: JY, JL, QX, QM, HS, SL, PD, and LL. Bioinformatic analyses, JL, SL, XZ, and CC, Writing, JY, JL, CC, QM, and MPM. Funding acquisition: XZ and PD. JY, JL, and QX, are joint first authors. All authors read and approved the manuscript.

## ACKNOWLEDGMENTS

This work was funded by the Natural Science Foundation of China (No. 82073159) and and the Science, Technology, and Innovation Commission of Shenzhen Municipality under the grant (No. JSGG20180508152912700).

## Competing interests

The authors declare that they have no competing interests.

## Notes

### Competing Interest Statement

The authors have declared no competing interest.

