## Supplementary figures and images for "Multi-omics analyses of single cell-derived colorectal cancer organoids reveal intratumor heterogeneity and immune response diversity"

### Supplemental Figure 1

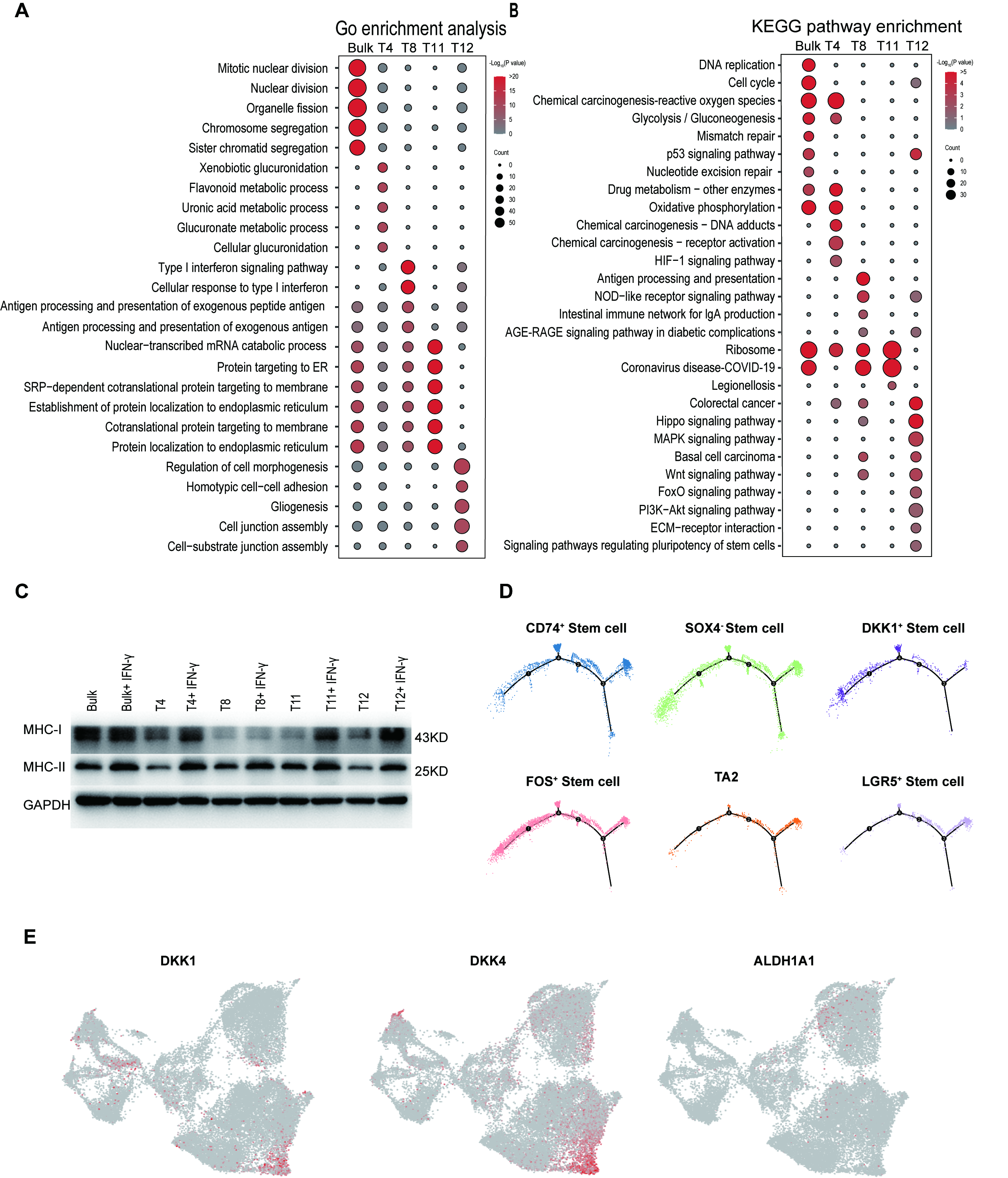

### Supplemental Figure 2

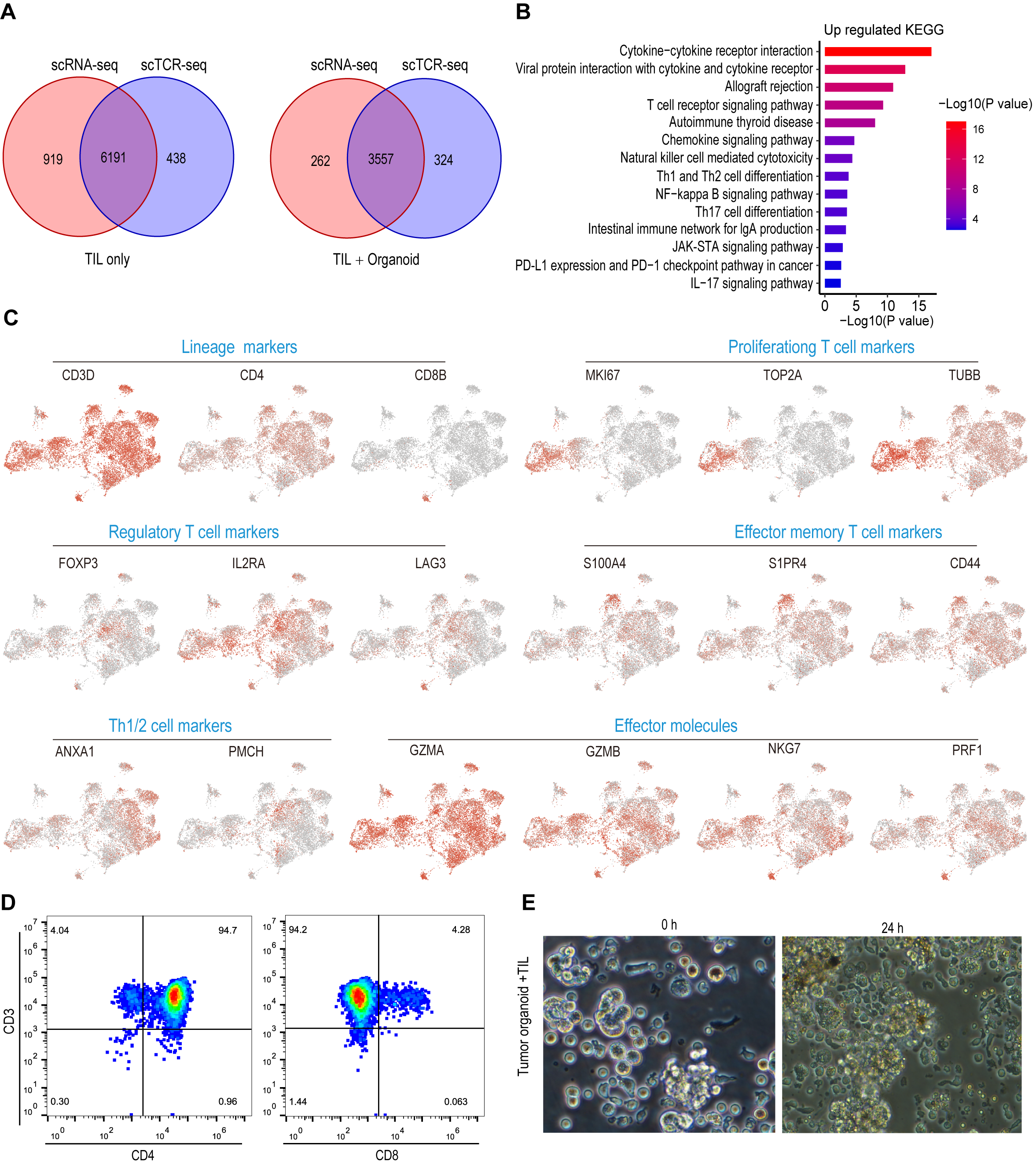
